# LOMDA: Linear optimization for miRNA-disease association prediction

**DOI:** 10.1101/751651

**Authors:** Yan-Li Lee, Ratha Pech, Maryna Po, Dong Hao, Tao Zhou

## Abstract

MicroRNAs (miRNAs) have been playing a crucial role in many important biological processes e.g., pathogenesis of diseases. Currently, the validated associations between miRNAs and diseases are insufficient comparing to the hidden associations. Testing all these hidden associations by biological experiments is expensive, laborious, and time consuming. Therefore, computationally inferring hidden associations from biological datasets for further laboratory experiments has attracted increasing interests from different communities ranging from biological to computational science. In this work, we propose an effective and efficient method to predict associations between miRNAs and diseases, namely linear optimization (LOMDA). The proposed method uses the heterogenous matrix incorporating of miRNA functional similarity information, disease similarity information and known miRNA-disease associations. Compared with the other methods, LOMDA performs best in terms of AUC (0.970), precision (0.566), and accuracy (0.971) in average over 15 diseases in local 5-fold cross-validation. Moreover, LOMDA has also been applied to two types of case studies. In the first case study, 30 predictions from breast neoplasms, 24 from colon neoplasms, and 26 from kidney neoplasms among top 30 predicted miRNAs are confirmed. In the second case study, for new diseases without any known associations, top 30 predictions from hepatocellular carcinoma and 29 from lung neoplasms among top 30 predicted miRNAs are confirmed.

**Author summary:** Identifying associations between miRNAs and diseases is significant in investigation of pathogenesis, diagnosis, treatment and preventions of related diseases. Employing computational methods to predict the hidden associations based on known associations and focus on those predicted associations can sharply reduce the experimental costs. We developed a computational method LOMDA based on the linear optimization technique to predict the hidden associations. In addition to the observed associations, LOMDA also can employ the auxiliary information (diseases and miRNAs similarity information) flexibly and effectively. Numerical experiments on global 5-fold cross validation show that the use of the auxiliary information can greatly improve the prediction performance. Meanwhile, the result on local 5-fold cross validation shows that LOMDA performs best among the seven related methods. We further test the prediction performance of LOMDA for two types of diseases based on HDMMv2.0 (2014), including (i) diseases with all the known associations, and (ii) new diseases without known associations. Three independent or updated databases (dbDEMC, 2010; miR2Disease, 2009; HDMMv3.2, 2019) are introduced to evaluate the prediction results. As a result, most miRNAs for target diseases are confirmed by at least one of the three databases. So, we believe that LOMDA can guide experiments to identify the hidden miRNA-disease associations.

## Introduction

MicroRNAs (miRNAs) are short for non-coding RNAs about 22 nucleotides that regulate gene expression of target post-transcriptional level [1–4]. In the last few decades, accumulative evidences show that miRNAs have strong relationships with many critical life processes including early cell growth, proliferation, apoptosis, differentiation and metabolism [5–9]. Moreover, miRNA dysregulation has also been shown to have close relation with many human complex diseases [10–14], including lung cancer [15] ,breast cancer [16, 17], cardiovascular diseases [18], and so on. Therefore, studying associations of miRNAs and diseases from biological datasets has become a significant problem in biomedical research which not only helps in investigation of pathogenesis [19], but also assists diagnosis, treatment, and preventions [20–23]. Biological experiments to verify new associations one by one would require a huge amount of time and labor, hence, an effective and efficient tool for selecting a small portion which are the most likely associations among a large pool of associations is needed for scientists to further experiments.

To investigate the miRNA-disease associations, comprehensive databases about miRNA-disease associations have been constructed, e.g., human miRNA disease database (HMDDv2.0) [24] collecting human miRNAs and diseases associations which are experiment-supported, dbDEMC [25] containing different expressions of miRNAs in human cancers detected by high-throughput methods, and miR2Disease [26] containing comprehensive resource of miRNA deregulations in various human diseases. These databases have facilitated researchers and scientists in understanding disease pathogenesis, furthermore, they are the main resources for association identification. Although there are rich collections about miRNA-disease association databases, these known associations are still limited comparing to all hidden miRNA-disease associations. Moreover, it is believed that one miRNA can be associated with multiple diseases and vice versa.

There is plenty of research [27] in predicting associations between miRNAs and diseases by using computational methods [28–32] and network-based methods [33–39]. Specifically, Chen *et al*. [34] proposed a method named random walk with restart for miRNA–disease associations (RWRMDA) to predict novel human miRNA-disease associations. In their proposed method, two datasets are used including the miRNA-disease network and miRNA functional similarities. Random walk cannot reach nodes that are isolated from the others. Therefore, it cannot predict associations between miRNAs and diseases that do not have any known associations. Similarly, Xuan *et al*. [40] proposed a method namely MIDP (short for miRNAs associated with Diseases Prediction), which is also based on the random walk technique, and the miRNA-disease network and miRNA functional similarities are used in their model. The difference between RWRMDA and MIDP is that the transition matrices are different. Chen *et al*. [41] developed a method by using the semi-supervised learning technique namely the regularized least squares for miRNA-disease association (RLSMDA) prediction, which integrates miRNA-disease associations, disease-disease similarities, and miRNA-miRNA similarities. Treating association prediction problem as binary classification would suffer from one drawback such that known associations are used as positive labels, while some unknown associations are used as negative labels. However, these unknown associations could be the hidden associations which are actually with positive labels. Zeng *et al.* [42] proposed a method by using structural perturbation method (SPM) on the integration of miRNA-disease association network, miRNA similarity network, and disease similarity network. Zeng *et al.* [43] also proposed a method namely neural network model for miRNA-disease association prediction (NNMDA), which also uses heterogenous network by integrating neighborhood information. SPM and NNMDA can well integrate multiple biological data resources to do miRNAs-diseases association prediction.

In this work, we propose an effective and efficient method namely linear optimization for miRNA-disease associations prediction (LOMDA) to infer associations between miRNAs and diseases in both cases: either (i) diseases and miRNAs similarity information is available, or (ii) only known associations are provided. LOMDA can also predict associations between miRNAs and diseases that do not have any known associations, but have miRNAs functional similarity information or disease similarity information, respectively. The performance of LOMDA measured by AUC, precision, and accuracy has been shown much higher than other benchmarks. Moreover, case studies demonstrate the effectiveness of LOMDA on predicting novel associations between miRNAs and diseases. Results show that LOMDA would be a promising bioinformatics tool for biomedical researchers. The Matlab source code of the proposed method and datasets used in this work can be obtained at https://github.com/rathapech/LOMDA.

## Materials and methods

### Data description

The miRNA-disease associations and two types of auxiliary information will be introduced in this subsection.

The human miRNA–disease associations are obtained from HMDDv2.0 [24]. If a miRNA is related to a disease, there is one link between them. After removing duplications, there are 6313 associations between 577 miRNAs and 336 diseases. The corresponding adjacency matrix is denoted as **MD**.

The disease-disease similarity matrix is constructed based on the gene functional information. Genes with similar function regulate similar diseases with greater probability [37]. The functional similarity of genes here is characterized by the log-likelihood score (LLS) [44], which can be downloaded from the HumanNet database [45]. Furthermore, the similarity of diseases *d*_*i*_ and *d*_*j*_ is calculated as

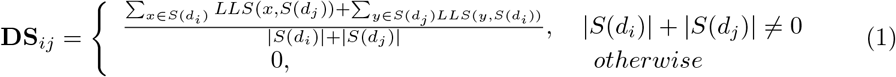

where *S*(*d*_*i*_) and *S*(*d*_*j*_) are gene sets that related to disease *d*_*i*_ and *d*_*j*_, respectively. |*S*| is the cardinality of set *S*. *LLS*(*x, S*(*d*_*i*_)) is *LLS* between the gene *x* and the gene set *S*(*d*_*i*_), where *x* ∈ *S*(*d*_*j*_). There are 56010 links between 336 diseases, and they can be formulated as a symmetric weighted matrix **DS**.

The miRNA-miRNA functional similarity matrix is obtained from four sources of information, including verified miRNA-target associations (*RST*), which can be obtained from miRTarBase [46], miRNA family information (*RSF* , ftp://mirbase.org/pub/mirbase/CURRENT/mi-Fam.dat.gz), cluster information (*RSC*, http://www.mirbase.org/) and verified miRNA-disease associations (*RSD*, http://www.cuilab.cn/files/images/cuilab/misim.zip) [37]. *RST*_*ij*_ is the number of shared targets between miRNAs *r*_*i*_ and *r*_*j*_. *RSF*_*ij*_ = 1 if two miRNAs belong to the same miRNA family, and *RSF*_*ij*_ = 0, otherwise. Similarly, *RSC*_*ij*_ = 1 if two miRNAs belong to the same cluster, and *RSC*_*ij*_ = 0, otherwise. RSD is calculated by MISIM (miRNA similarity) method [37]. Finally, **MS**(*r*_*i*_, *r*_*j*_) reads

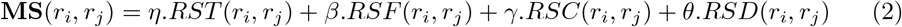

where *η, β, γ* and *θ* are parameters to adjust the four weights and are set as *η* = 0.2, *β* = 0.1, *γ* = 0.2 and *θ* = 0.5 [42]. There are 35521 links between 577 miRNAs, and they can be formulated as a symmetric weighted matrix **MS**.

The above three matrices can be combined to get an heterogenous matrix as

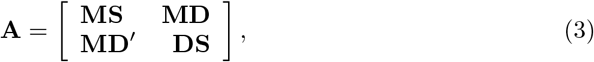

where **MS** ∈ ℝ^*m*×*m*^ is the miRNA functional similarity matrix in which *m* is the number of miRNAs; **DS** ∈ ℝ^*d*×*d*^ is the disease similarity matrix in which *d* is the number of diseases; **MD** ∈ ℝ^*m*×*d*^ is the known miRNA-disease association matrix. If only associations between miRNAs and diseases are available, we can set **A** = **MD**. The composition of heterogenous matrix **A** is shown in Fig 1.

**Fig 1.**
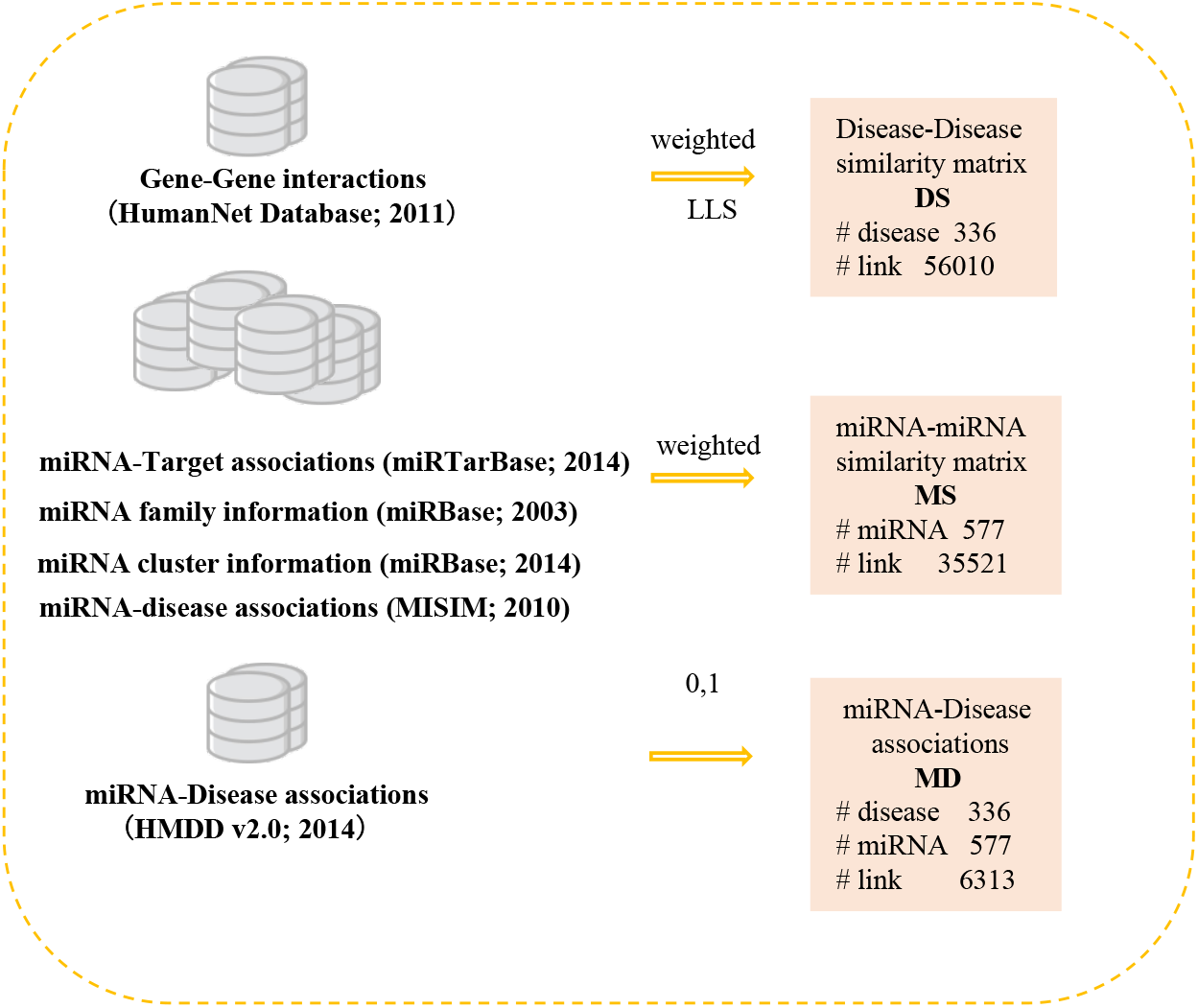
The illustration of the input datasets of LOMDA.

## LOMDA

Denoting the integration matrix by **A** as shown in Eq (3), we assume that the likelihood of associations between miRNAs and diseases can be written as a linear combination of **A** and weighting matrix **Z** as

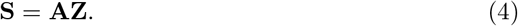

Since **S** and **Z** are unknown, the problem of Eq (4) has infinite solutions. However, in order to obtain the likelihood **S** containing existing and predicted associations, **S** should be intuitively and reasonably close to **A**. Then we can write

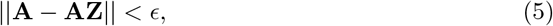

where *ϵ* is the threshold parameter. Moreover, to avoid the model to be overfitted and simultaneously to constrain the magnitudes of **Z** , we can relax the Eq (5) as

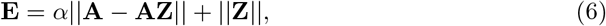

where *α* is the positive free parameter greater than 0 and ∥.∥ is the matrix norm. Without losing the generality [47], we use Frobenius norm and raise the two terms with power 2. We can have

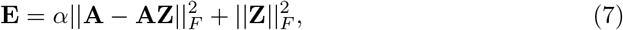

where Frobenius norm is denoted as 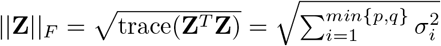, e.g. *σ*_*i*_ is the singular value, *p* is the number of row, and *q* is the number of column of **Z** . The expansion of Eq (7) reads

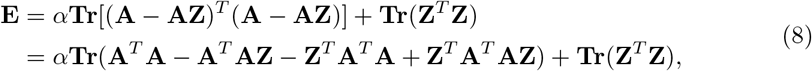

with its partial derivative being

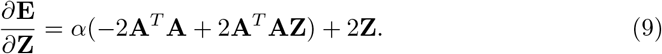

Setting *∂***E**/*∂***Z** = 0, we can obtain the optimal solution of **Z** as

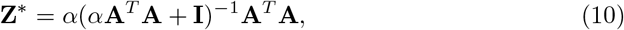

where **I** is the identity matrix. The likelihood matrix **S** can be obtained as

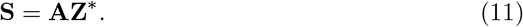

Finally, we can obtain scores of unobserved miRNAs and diseases associations based on **S**.

In addition, when only known association information of miRNAs and diseases is available, LOMDA can also effectively predict missing associations by using only **MD**. We replace **A** by matrix **MD** in Eq (4) through Eq (11) and keep the derivation the same. When only matrix **MD** is used, we call the proposed method as LOMDA-MD. We illustrate the performance of these two derived methods in result section.

## Results

### Performance evaluation

To evaluate our proposed method against others, we adopt the 5-fold cross validation technique to test the prediction performance. Moreover, two different prediction tasks are considered: (i) predicting hidden associations for all the diseases simultaneously. Specifically, all the associations in **MD** are divided into 5 disjoint subsets. Four subsets are treated as training samples, and the remaining subset is treated as testing samples.

We call it as global 5-fold cross validation; (ii) predicting hidden associations for some specific diseases, i.e., for a specific disease, the related associations in matrix **MD** are divided into 5 disjoint subsets. One subset is considered as the testing set, and the remaining associations in **MD** are considered as training set. We call it as local 5-fold cross validation. For the two tasks, we repeatedly and independently do the simulation for five times until all five subsets are used as testing samples exactly once. This 5-fold cross validation is repeated by 20 times. That is to say, the final evaluation metrics introduced in the following are the average over 20*5 prediction results.

We introduce three evaluation metrics including AUC, precision, and accuracy to evaluate the prediction performance. AUC can be interpreted as a probability that a randomly chosen hidden association (i.e., associations in testing samples) is given a higher score than a randomly chosen unknown association (i.e., associations that have not been collected in training samples and testing samples) [48]. Notice that, in the second task for a specific disease, unknown associations denote associations between the target disease and miRNAs that have not been collected by the target disease according to the matrix **MD**. We randomly choose *n* pairs of associations from the testing samples and the unknown associations, respectively. *n*_1_ denotes the number of pairs with higher score for hidden associations, and *n*_2_ denotes the number of pairs with the same score between the hidden association and the unknown association. The score values are obtained based on the related method. Thus, 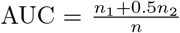 , and *n* is set to 10000 in this work. Precision is defined as the ratio of right predicted associations (i.e., true positive associations) to all the predicted associations. We select top *L* associations in predicted scores and count them if they exist in testing data, and the hitting number is denoted as *L*_*r*_, then 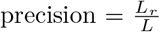. *L* is the number of testing samples in our work. Accuracy is obtained by the ratio of true positive associations and true negative associations to all the candidate associations, i.e., the union set of hidden associations and unknown associations.

### Performance with auxiliary information

In this subsection, LOMDA and LOMDA-MD are used to test the contribution of auxiliary information to prediction results. The prediction is conducted for all the diseases simultaneously by the global 5-fold cross validation technique. For LOMDA, not only the training set but also the auxiliary information **MS** and **DS** are employed to do prediction, while for LOMDA-MD, only the training set is used. As shown in Fig 2, LOMDA performs much better than LOMDA-MD. Meanwhile, we can find that LOMDA achieves its best performance on *α* = 0.01, and *α* = 0.001 for LOMDA-MD.

**Fig 2.**
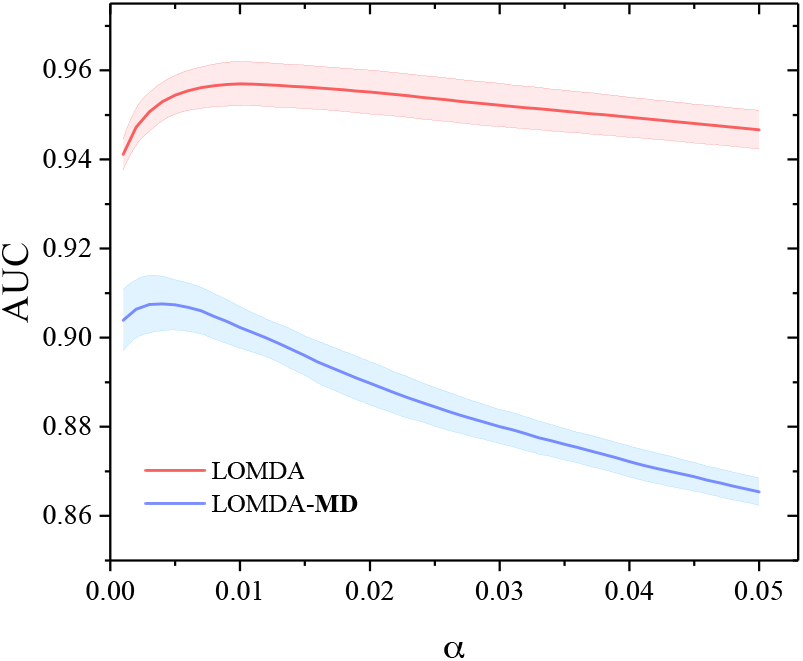
The contribution of auxiliary information to prediction results based on global 5-fold cross validation. The shaded area presents the standard deviation.

### Comparison with other methods

A great number of methods are developed to predict hidden associations between miRNAs and diseases, including RWRMDA [34], RLSMDA [41], MIDP [40], SPM [42] and NNMDA [43]. Detailed information employed by each method is shown in Table 1. The auxiliary information (**MS** and **DS**) are employed according to Table 1. The prediction is conducted for some specific diseases based on local 5-fold cross validation. Table 2–Table 4 illustrate the AUC, precision, and accuracy of the proposed method and others on 15 different diseases, respectively. The highest values generating from any methods are shown in boldface. As shown in the bottom row of each table, average AUC values among the 15 diseases of RWRMDA, RLSMDA, MIDP, SPM, NNMDA, LOMDA-MD and LOMDA are 0.898, 0.950, 0.953, 0.786, 0.920, 0.938, 0.970, respectively. Average precision values on all the 15 diseases obtained from RWRMDA, RLSMDA, MIDP, SPM, NNMDA, LOMDA-MD and LOMDA are 0.207, 0.417, 0.392, 0.306, 0.442, 0.372 and 0.566, respectively. Average accuracy values on the 15 diseases from RWRMDA, RLSMDA, MIDP, SPM, NNMDA, LOMDA-MD and LOMDA are 0.944, 0.961, 0.959, 0.950, 0.938, 0.956 and 0.971, respectively. Overall speaking, LOMDA outperforms the other benchmarks on AUC, precision and accuracy.

**Table 1.** Information included in seven methods. **MS** is the miRNA-miRNA functional similarity matrix, **DS** is the disease-disease similarity matrix, and **MD** is the known miRNA-disease association matrix.

| Datasets | RWRMDA | RLSMDA | MIDP | SPM | NNMDA | LOMDA-MD | LOMDA |
| --- | --- | --- | --- | --- | --- | --- | --- |
| **MS** | Yes | Yes | Yes | Yes | Yes | No | Yes |
| **DS** | No | Yes | No | Yes | Yes | No | Yes |
| **MD** | Yes | Yes | Yes | Yes | Yes | Yes | Yes |

**Table 2.**
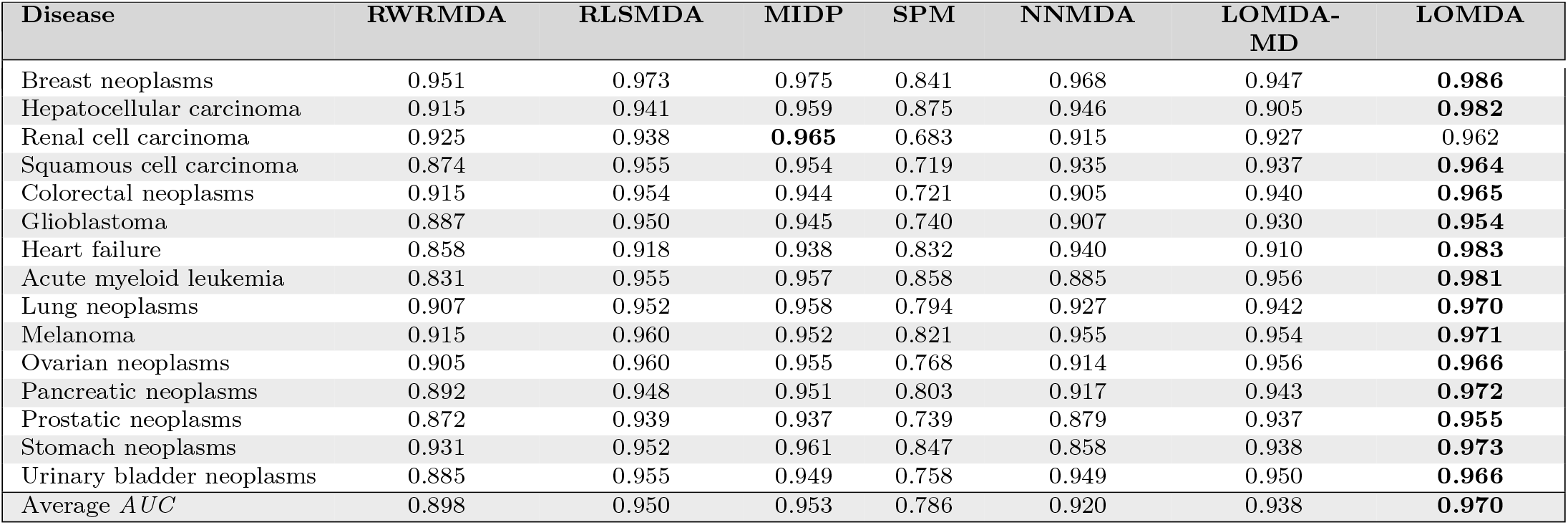
AUC of different methods by using local 5-fold cross validation on 15 diseases.

**Table 3.**
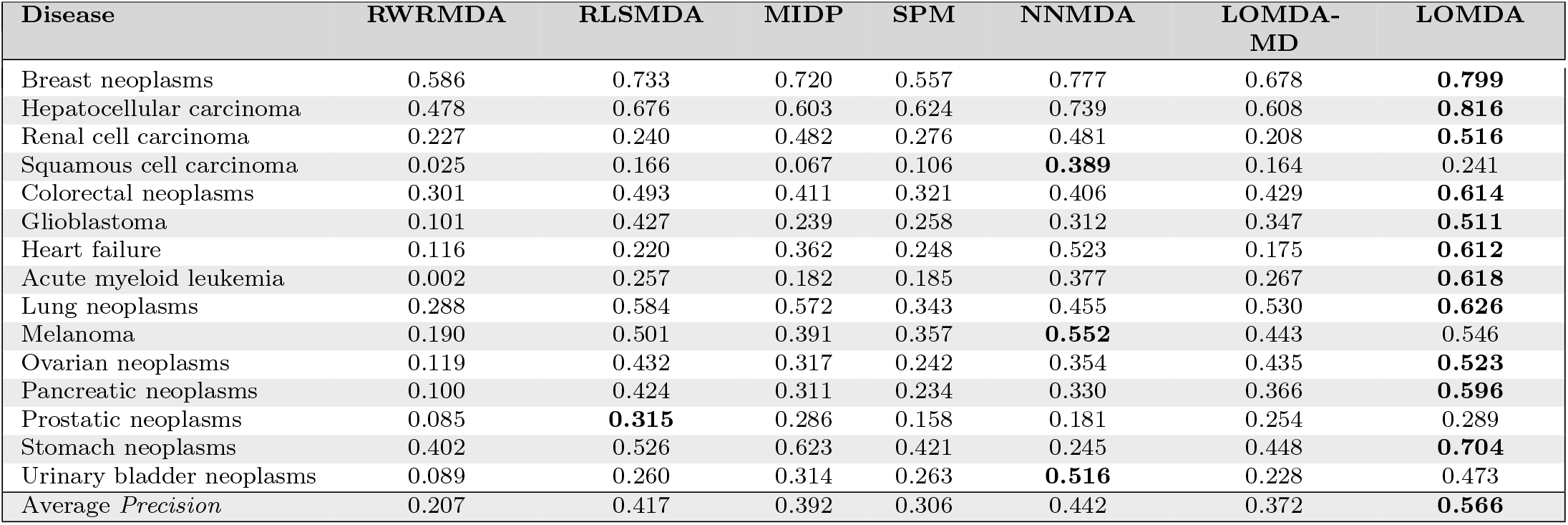
Precision of different methods by using local 5-fold cross validation on 15 diseases.

| Disease | RWRMDA | RLSMDA | MIDP | SPM | NNMDA | LOMDA-MD | LOMDA |
| --- | --- | --- | --- | --- | --- | --- | --- |
| Breast neoplasms | 0.586 | 0.733 | 0.720 | 0.557 | 0.777 | 0.678 | **0.799** |
| Hepatocellular carcinoma | 0.478 | 0.676 | 0.603 | 0.624 | 0.739 | 0.608 | **0.816** |
| Renal cell carcinoma | 0.227 | 0.240 | 0.482 | 0.276 | 0.481 | 0.208 | **0.516** |
| Squamous cell carcinoma | 0.025 | 0.166 | 0.067 | 0.106 | **0.389** | 0.164 | 0.241 |
| Colorectal neoplasms | 0.301 | 0.493 | 0.411 | 0.321 | 0.406 | 0.429 | **0.614** |
| Glioblastoma | 0.101 | 0.427 | 0.239 | 0.258 | 0.312 | 0.347 | **0.511** |
| Heart failure | 0.116 | 0.220 | 0.362 | 0.248 | 0.523 | 0.175 | **0.612** |
| Acute myeloid leukemia | 0.002 | 0.257 | 0.182 | 0.185 | 0.377 | 0.267 | **0.618** |
| Lung neoplasms | 0.288 | 0.584 | 0.572 | 0.343 | 0.455 | 0.530 | **0.626** |
| Melanoma | 0.190 | 0.501 | 0.391 | 0.357 | **0.552** | 0.443 | 0.546 |
| Ovarian neoplasms | 0.119 | 0.432 | 0.317 | 0.242 | 0.354 | 0.435 | **0.523** |
| Pancreatic neoplasms | 0.100 | 0.424 | 0.311 | 0.234 | 0.330 | 0.366 | **0.596** |
| Prostatic neoplasms | 0.085 | **0.315** | 0.286 | 0.158 | 0.181 | 0.254 | 0.289 |
| Stomach neoplasms | 0.402 | 0.526 | 0.623 | 0.421 | 0.245 | 0.448 | **0.704** |
| Urinary bladder neoplasms | 0.089 | 0.260 | 0.314 | 0.263 | **0.516** | 0.228 | 0.473 |
| Average *Precision* | 0.207 | 0.417 | 0.392 | 0.306 | 0.442 | 0.372 | **0.566** |

**Table 4.** Accuracy of different methods by using local 5-fold cross validation on 15 diseases.

| Disease | RWRMDA | RLSMDA | MIDP | SPM | NNMDA | LOMDA-MD | LOMDA |
| --- | --- | --- | --- | --- | --- | --- | --- |
| Breast neoplasms | 0.926 | 0.951 | 0.949 | 0.919 | 0.943 | 0.941 | **0.963** |
| Hepatocellular carcinoma | 0.900 | 0.938 | 0.925 | 0.928 | 0.932 | 0.925 | **0.965** |
| Renal cell carcinoma | 0.961 | 0.962 | 0.975 | 0.964 | 0.942 | 0.961 | **0.976** |
| Squamous cell carcinoma | 0.976 | 0.980 | 0.976 | 0.977 | 0.956 | 0.980 | **0.981** |
| Colorectal neoplasms | 0.931 | 0.950 | 0.941 | 0.932 | 0.912 | 0.944 | **0.962** |
| Glioblastoma | 0.955 | 0.971 | 0.964 | 0.963 | 0.939 | 0.967 | **0.976** |
| Heart failure | 0.948 | 0.954 | 0.963 | 0.955 | 0.947 | 0.952 | **0.977** |
| Acute myeloid leukemia | 0.974 | 0.981 | 0.979 | 0.979 | 0.967 | 0.981 | **0.990** |
| Lung neoplasms | 0.931 | 0.960 | 0.959 | 0.937 | 0.932 | 0.955 | **0.964** |
| Melanoma | 0.928 | 0.956 | 0.945 | 0.942 | 0.933 | 0.950 | **0.959** |
| Ovarian neoplasms | 0.951 | 0.969 | 0.963 | 0.958 | 0.940 | 0.967 | **0.972** |
| Pancreatic neoplasms | 0.946 | 0.965 | 0.961 | 0.954 | 0.942 | 0.961 | **0.976** |
| Prostatic neoplasms | 0.946 | **0.960** | 0.958 | 0.949 | 0.921 | 0.958 | **0.960** |
| Stomach neoplasms | 0.923 | 0.940 | 0.951 | 0.924 | 0.903 | 0.929 | **0.961** |
| Urinary bladder neoplasms | 0.967 | 0.973 | 0.976 | 0.973 | 0.962 | 0.971 | **0.980** |
| Average *AUC* | 0.944 | 0.961 | 0.959 | 0.950 | 0.938 | 0.956 | **0.971** |

### Case studies

In this subsection, two detailed case studies for five critical diseases including breast neoplasms, colon neoplasms, kidney neoplasms, hepatocellular carcinoma and lung cancer have been investigated by LOMDA. These five diseases have attracted widespread attention from the general public and academia. Breast neoplasms is the most common malignant tumor in women accounting for 25% [49] followed by prostate and colon cancer [50, 51]. Colon neoplasm is one of the common cancers which has high death rate [52]. The incidence of kidney cancer has increased 43% since 1973, and the risk of the disorder is higher in men than in women and increases with age [53]. Hepatocellular cancer is the third leading cause of cancer-related deaths worldwide [54]. Lung cancer is till a leading cause of cancer death in both men and women in the United States even though its extensive list of risk factors has been characterized [55].

In the first case study, we test the performance of LOMDA in predicting new associated miRNAs for breast neoplasms, colon neoplasms and kidney neoplasms based on **MD** (HMDDv2.0; 2014), **DS** (HumanNet database; 2011) and **MS** (miRTarBase; 2014, miRBase; 2003, MISIM; 2010). All the known associations in **MD** together with all the similarity information **MS** and **DS** are utilized to do prediction. Three independent or updated databases are introduced to verify the the prediction results, including dbDEMC (2010) [25], miR2Disease (2009) [26] and HMDDv3.2 (2019) [56]. First of all, for the target disease, we compute scores for all the candidate associations then sort them in descending order. After that, we select the top 30 association scores of the interested disease and manually verify the existences of the associations by the above three databases. Results for three diseases including breast neoplasms, colon neoplasms, and kidney neoplasms are shown in Table 5, Table 6, and Table 7, respectively. We can find that 30 out of the top 30 predicted breast neoplasms-related miRNAs are confirmed, 24 out of the top 30 predicted colon neoplasms-related miRNAs are confirmed, and 26 of the top 30 predicted kidney neoplasms-related miRNAs are confirmed.

**Table 5.** The top 30 predicted miRNAs associated with breast neoplasms. The first column records top 1-15 related miRNAs, and the third column records top 16-30 miRNAs.

| miRNA | Evidence | miRNA | Evidence |
| --- | --- | --- | --- |
| hsa-mir-106a | dbDEMC; HMDDv3.2 | hsa-mir-138 | dbDEMC; HMDDv3.2 |
| hsa-mir-142 | HMDDv3.2 | hsa-mir-574 | HMDDv3.2 |
| hsa-mir-130a | dbDEMC; HMDDv3.2 | hsa-mir-15b | dbDEMC; HMDDv3.2 |
| hsa-mir-99a | dbDEMC; HMDDv3.2 | hsa-mir-19b | dbDEMC; HMDDv3.2 |
| hsa-mir-150 | dbDEMC; HMDDv3.2 | hsa-mir-30e | HMDDv3.2 |
| hsa-mir-378a | HMDDv3.2 | hsa-mir-542 | HMDDv3.2 |
| hsa-mir-186 | dbDEMC | hsa-mir-650 | dbDEMC |
| hsa-mir-92b | dbDEMC; HMDDv3.2 | hsa-mir-98 | dbDEMC; miR2Disease; HMDDv3.2 |
| hsa-mir-185 | dbDEMC; HMDDv3.2 | hsa-mir-370 | dbDEMC; HMDDv3.2 |
| hsa-mir-130b | dbDEMC; HMDDv3.2 | hsa-mir-99b | dbDEMC |
| hsa-mir-192 | dbDEMC; HMDDv3.2 | hsa-mir-95 | dbDEMC |
| hsa-mir-212 | dbDEMC; HMDDv3.2 | hsa-mir-32 | dbDEMC; HMDDv3.2 |
| hsa-mir-449b | dbDEMC | hsa-mir-196b | dbDEMC; HMDDv3.2 |
| hsa-mir-372 | dbDEMC; HMDDv3.2 | hsa-mir-449a | dbDEMC; HMDDv3.2 |
| hsa-mir-330 | dbDEMC | hsa-mir-181c | dbDEMC; HMDDv3.2 |

**Table 6.** The top 30 predicted miRNAs associated with colon neoplasms. The first column records top 1-15 related miRNAs, and the third column records top 16-30 miRNAs.

| miRNA | Evidence | miRNA | Evidence |
| --- | --- | --- | --- |
| hsa-mir-200a | dbDEMC; HMDDv3.2 | hsa-mir-224 | dbDEMC; HMDDv3.2 |
| hsa-mir-92a | dbDEMC; HMDDv3.2 | hsa-mir-429 | dbDEMC; HMDDv3.2 |
| hsa-mir-29b | dbDEMC; HMDDv3.2 | hsa-mir-34b | *unconfirmed* |
| hsa-mir-222 | dbDEMC; HMDDv3.2 | hsa-mir-199a | *unconfirmed* |
| hsa-mir-34c | *unconfirmed* | hsa-mir-148a | dbDEMC; HMDDv3.2 |
| hsa-mir-100 | dbDEMC | hsa-mir-27a | dbDEMC; miR2Disease; HMDDv3.2 |
| hsa-mir-210 | dbDEMC; HMDDv3.2 | hsa-mir-181b | dbDEMC; miR2Disease; HMDDv3.2 |
| hsa-mir-182 | dbDEMC | hsa-mir-195 | dbDEMC; HMDDv3.2 |
| hsa-mir-375 | dbDEMC; HMDDv3.2 | hsa-mir-20b | dbDEMC |
| hsa-mir-1-2 | dbDEMC | hsa-mir-150 | dbDEMC; HMDDv3.2 |
| hsa-mir-203 | dbDEMC; miR2Disease; HMDDv3.2 | hsa-mir-103a | *unconfirmed* |
| hsa-mir-181a | dbDEMC; miR2Disease; HMDDv3.2 | hsa-mir-181a-2 | *unconfirmed* |
| hsa-mir-30d | dbDEMC; HMDDv3.2 | hsa-mir-146b | dbDEMC |
| hsa-mir-99a | dbDEMC | hsa-mir-25 | dbDEMC; miR2Disease; HMDDv3.2 |
| hsa-mir-29c | dbDEMC | hsa-mir-151a | *unconfirmed* |

**Table 7.** The top 30 predicted miRNAs associated with kidney neoplasms. The first column records top 1-15 related miRNAs, and the third column records top 16-30 miRNAs.

| miRNA | Evidence | miRNA | Evidence |
| --- | --- | --- | --- |
| hsa-mir-21 | dbDEMC; miR2Disease; HMDDv3.2 | hsa-mir-34c | dbDEMC |
| hsa-mir-146a | dbDEMC | hsa-mir-30a | dbDEMC |
| hsa-mir-17 | dbDEMC; miR2Disease; HMDDv3.2 | hsa-mir-638 | dbDEMC |
| hsa-mir-200a | dbDEMC; miR2Disease; HMDDv3.2 | hsa-mir-320a | dbDEMC |
| hsa-mir-451a | dbDEMC | hsa-mir-499a | *unconfirmed* |
| hsa-mir-200b | dbDEMC; miR2Disease | hsa-mir-1207 | *unconfirmed* |
| hsa-mir-433 | *unconfirmed* | hsa-let-7b | dbDEMC |
| hsa-mir-192 | dbDEMC; HMDDv3.2 | hsa-mir-362 | dbDEMC; HMDDv3.2 |
| hsa-mir-198 | dbDEMC | hsa-mir-148b | *unconfirmed*; HMDDv3.2 |
| hsa-mir-205 | dbDEMC; miR2Disease | hsa-mir-199a | dbDEMC; miR2Disease; HMDDv3.2 |
| hsa-mir-423 | dbDEMC | hsa-mir-15a | dbDEMC; miR2Disease; HMDDv3.2 |
| hsa-mir-200c | dbDEMC; miR2Disease; HMDDv3.2 | hsa-mir-181a | dbDEMC |
| hsa-mir-30b | dbDEMC | hsa-mir-206 | dbDEMC |
| hsa-mir-10a | dbDEMC | hsa-mir-182 | dbDEMC; miR2Disease |
| hsa-mir-377 | dbDEMC | hsa-mir-196b | dbDEMC |

In the second case study, we verify the prediction performance of LOMDA on diseases without any known associations based on **MD** (HMDDv2.0; 2014), **DS** (HumanNet database; 2011) and **MS** (miRTarBase; 2014, miRBase; 2003, MISIM; 2010). Hepatocellular carcinoma and lung neoplasms are taken as examples. we remove all known associations of hepatocellular carcinoma and lung neoplasms from **MD** in **A**, respectively. Then we compute the likelihood scores of these diseases with all the miRNAs by using LOMDA. Finally, we select the top 30 candidates and manually check these candidates by the above three databases. All these predicted candidates belonging to hepatocellular carcinoma can be confirmed by at least one of the three databases and 29 among 30 predicted associations of lung neoplasms are also confirmed. The results are shown in Table 8 and Table 9, respectively.

**Table 8.**
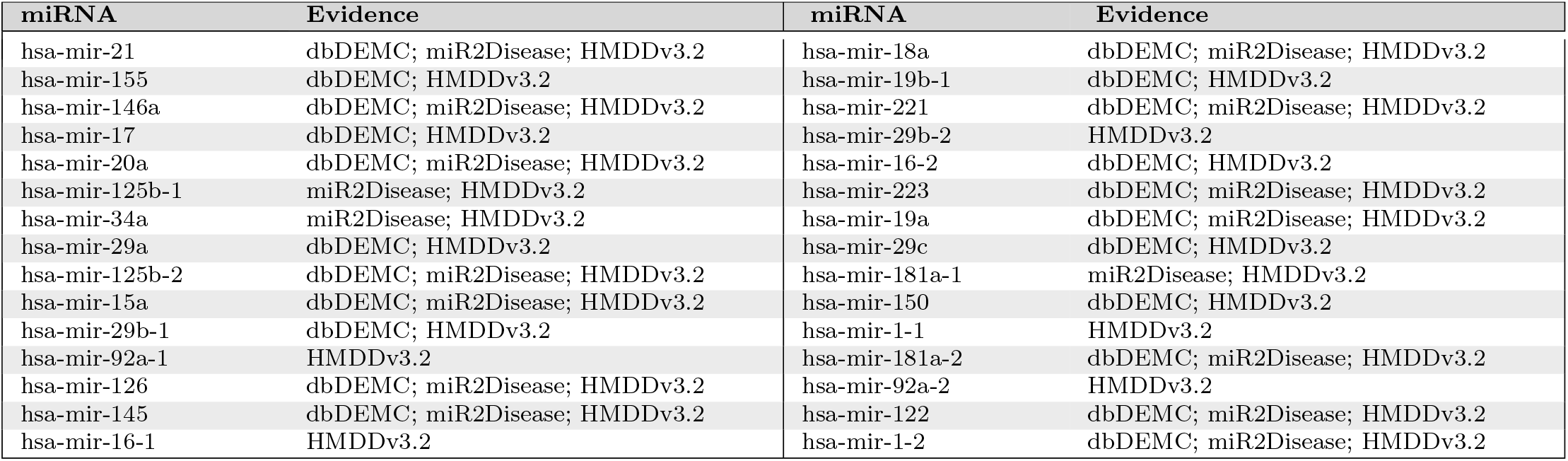
The top 30 miRNAs associated with hepatocellular carcinoma were predicted by LOMDA with hiding all known related miRNAs.

**Table 9.** The top 30 miRNAs associated with lung neoplasms were predicted by LOMDA with hiding all known related miRNAs.

| miRNA | Evidence | miRNA | Evidence |
| --- | --- | --- | --- |
| hsa-mir-21 | dbDEMC; miR2Disease; HMDDv3.2 | hsa-mir-146a | dbDEMC; miR2Disease; HMDDv3.2 |
| hsa-mir-34c | dbDEMC; HMDDv3.2 | hsa-mir-30e | dbDEMC; HMDDv3.2 |
| hsa-mir-155 | dbDEMC; HMDDv3.2 | hsa-mir-152 | dbDEMC; HMDDv3.2 |
| hsa-mir-486 | dbDEMC; HMDDv3.2 | hsa-mir-219-2 | *unconfirmed* |
| hsa-mir-15a | dbDEMC; HMDDv3.2 | hsa-mir-223 | HMDDv3.2 |
| hsa-mir-34a | dbDEMC; HMDDv3.2 | hsa-mir-520c | HMDDv3.2 |
| hsa-mir-148b | dbDEMC; HMDDv3.2 | hsa-mir-137 | dbDEMC; HMDDv3.2 |
| hsa-mir-126 | dbDEMC; HMDDv3.2 | hsa-mir-134 | dbDEMC |
| hsa-mir-29a | dbDEMC; miR2Disease; HMDDv3.2 | hsa-mir-125b | dbDEMC; miR2Disease; HMDDv3.2 |
| hsa-mir-200b | dbDEMC; miR2Disease; HMDDv3.2 | hsa-mir-204 | dbDEMC; HMDDv3.2 |
| hsa-mir-326 | dbDEMC; HMDDv3.2 | hsa-mir-374a | HMDDv3.2 |
| hsa-mir-34b | dbDEMC; HMDDv3.2 | hsa-mir-7b | miR2Disease; HMDDv3.2 |
| hsa-mir-337 | dbDEMC; HMDDv3.2 | hsa-mir-375 | dbDEMC; HMDDv3.2 |
| hsa-mir-25 | dbDEMC; HMDDv3.2 | hsa-mir-92b | dbDEMC; HMDDv3.2 |
| hsa-mir-122 | dbDEMC HMDDv3.2 | hsa-mir-28 | dbDEMC; HMDDv3.2 |

## Conclusion

Predicting novel associations between miRNAs and diseases helps scientists firstly focus on the most likely associations rather than blindly check on all possible associations which is extremely costly and laborious. Moreover, it can help researchers enhance their understanding toward molecular mechanisms of diseases at the miRNA level. This prediction also plays an important role in understanding the pathogenesis of human diseases at the early stage, therefore, it can help in diagnosis, treatment and prevention.

Motivated by the necessity of identifying novel associations between miRNAs and diseases, in this work we propose a computational method, namely linear optimization for miRNA-disease association (LOMDA) prediction. The proposed method utilizes the heterogenous matrix by integrating miRNA functional similarity information, disease similarity information, and known miRNA-disease associations. In case only known associations are available, the method can also be applied. Moreover, the method can also predict associations for new miRNAs (or diseases) by using miRNA functional similarity information (or disease similarity information). According to the cross validation evaluated by AUC, precision, and accuracy, the proposed method has been shown to perform very satisfactorily. Thus, LOMDA is an effective and efficient tool for predicting miRNA-disease associations.

## Acknowledgments

This work was partially supported by the National Natural Science Foundation of China (Grant Nos. 11975071 and 61703074).

